# It takes two vanilloid ligand bindings per channel to transduce painful capsaicin stimuli

**DOI:** 10.1101/725598

**Authors:** Ting-Yi Liu, Ying Chu, Hao-Ruei Mei, Dennis S. Chang, Huai-Hu Chuang

## Abstract

The capsaicin receptor TRPV1 in mammals evolved the capability of integrating pain arising from combinations of noxious temperature or chemical irritants. Four-fold repetition of TRPV1 subunits makes an ion channel endowed with excellent sensitivity for pain detection, assisting this ionotropic receptor to differentiate graded injuries. We manipulated the stoichiometry and relative steric coordination of capsaicin binding at the molecular level, explicating rules with which a receptor codes pain within a broad range of intensity. The first ligand binding delivers small but clear initiation of channel activation. Maximal agonist action has already been reached in a receptor-in-tandem containing two or three wild-type receptor units, displaying activity comparable to the full liganded all-wild-type tandem tetramers. When the binding sites outnumbered ligands, independent action dominates in each channel. The non-vanilloid agonist 2-APB differs from capsaicin by adopting a distinct open mechanism since it does not demand a vanilloid group to activate. The sharing of the same pore greatly simplifies synergism to transduce relevant inputs by summation for pain signaling. And questions the need to explore deeper into other aspects of nociception.

## Introduction

Transient receptor potential vanilloid (TRPV1) is a multi-subunit cation-permeating channel assembled from four identical subunits jointly surrounding the ion-conducting pore (Caterina et al., 1997; Clapham D.E. 2003; Liao et al., 2013). TRPV1 responds to both physical challenge like noxious heat and harmful environmental chemical (protons, irritants, and inflammatory mediators) (Caterina et al., 1997, 2000; Davis et al., 2000; Macpherson et l., 2005; Nieto-Posadas et al., 2011; Siemens et al., 2006; Tominaga et al., 1998), converting them into electrical activity in sensory neurons for chemical or thermal nociception (Baron et al., 2006; Julius et al., 2013; Marrone et al., 2017).

Built for detecting imminent tissue injury, TRPV1 must be sensitive enough to detect the insult and deliver a timely response. Besides, it obliges just sufficient sensitivity for sensible decoding of external or internal inflicts about to compromise the animal’s physiological capacity. Nature designs TRPV1 with multiplicity of same repeats, making it a sensitive signal transduction apparatus. TRPV1 thus provides a substrate for creating a complex of the potential to broadly quantify combinations of environmental biochemical events. In particular, micro-fluorometry was employed frequently to quantify intracellular Ca^2+^ rise from ionic influx through rTRPV1.

Capsaicin is a powerful ligand for TRPV1; it brings about cation influx, primary sensory neuronal action potentials and typically a burning sensation in mammals (Caterina et al., 1997, Wood et al., 1988). “Computational” integration, within neurons dictates the ultimate TRPV1 response amplitudes subsequent to signal convergence (Hazan et al., 2015), of which the output relates well to capsaicin-provoked chemical nociception. S512F, essentially a loss-of-vanilloid binding mutant subunit (Jordet S.E. and Julius D. 2002), was mixed with wild type to tune the capacity of ligand induced channel opening. The ability of a TRPV1 mutant to relay capsaicin excitation is dictated by the number of S512F subunits in a channel complex.

Notwithstanding the creation any combinations of point mutations, analysis of receptor activation is also complicated by heterogeneity of compositional stoichiometry upon subunit assemblies (Chen et al., 2013; Robinson and Sauer R.T. 1998). We surmounted this complication by constructing tandem tetrameric receptors; the number of the wild-type subunits thus became exacted, vastly simplifying the mechanistic analyses of activation of TRPV1 tetramers with distinct arrangements. Above all, functional expressions of tandem receptors expressed properly as channels assembled from monomeric subunits. Multiple capsaicin-bound subunits are required to drive the channel to maximal opening.

## Materials and Methods

### Molecular biology - multimeric rTRPV1 constructs

The QuikChange site-directed mutagenesis was used to get the single point mutants by overlap extension PCR with Phusion polymerase (New England Biolab). Tetramers were constructed by linking four TRPV1 genes with the inter-subunit hepta-peptide linker ANENGDA between N-terminus and C-terminus followed by restriction digestion and ligation (Chen et al., 2013; Robinson C.R. and Sauer R.T. 1998)

### Cell culture and heterologous expression

Human embryonic kidney 293 T (HEK293T) cells were maintained in MEM/EBSS (HyClone) supplemented with 10% fetal bovine serum (FBS, Gibco), 100U/ml penicillin and 100μg/ml streptomycin (Lonza). HEK293T cells were transfected with 1-2 μg wild-type, mutant or tetramers receptor plasmids using Avalanche-Omni transfection reagent (EZ Biosystems). Cells were reseeded on 96-well coated with poly-D-lysine with (0.1 mg/ml) and collagen (55 μg/ml) from 24-36 hours after transfection, and then we did the calcium-imaging assay on the next day.

### Ratiometric calcium imaging

All calcium imaging experiments were conducted at 22°C. Cells were loaded with Ca^2+^ indicator Fura-2, AM or Fura-4F, AM (2 μM, Thermo Fisher Scientific) in 1.7-fold OR-2 solution (8.5mM HEPES, 140.3mM NaCl, 3.4mM KCl, 1.7mM MgCl_2_ and 1mM CaCl_2_, pH 7.4), the calcium replacement buffer with 1mM/10mM BaCl_2_ or 1mM/10mM SrCl_2_, or the sodium to cesium replacement buffer made of 140.3mM CsCl; all of the samples were incubated at 30°C for 3 hours, and then cells were washed with bath with 1mM EGTA solution away after Ca^2+^ indicator precursor Fura-AM removal and changed to the recording bath. Fluorescence data were acquired by capturing the frame rate at one frame every 2 sec (to record capsaicin dose response) or 5 sec (to record channel sensitization by H_2_O_2_), with 20-50 ms exposure time to either wavelength (340 and 380 nm) for excitation using an EMCCD camera (Photometrics, Evolve) driven by the Slidebook 6 digital microscopy software (Intelligent Imaging Innovations). The ± sign in this report is a standard error (SEM) calculated with sample number listed in figure legends and tables.

### Non-reducing SDS-PAGE analysis and Western blotting

Following the transfection, HEK293T cells were lysed with lysis buffer containing 0.5% Triton X-100, 0.5% NP-40 and protease inhibitor. To test the reversibility of covalent bonds, the cell lysates were mixed with 6× SDS non-reducing sample buffer or 6× SDS reducing sample buffer containing 2% dithiothreitol and 5% 2-mercaptoethanol. The lysates were resolved by 7.5% non-reducing SDS-PAGE, and the proteins were transferred onto a PVDF Transfer Membrane (Millipore). The membrane was incubated in blocking buffer (5% nonfat milk in Tris-buffered saline with 0.05% Tween 20) containing anti-rat TRPV1 antibody (GeneTex) at 1:5000 dilution or anti-GAPDH antibody (Santa Cruz Biotechnology) at 1:5000 dilution. Both of the proteins were visualized using a secondary anti-rabbit HRP-conjugated antibody (Thermo Fisher Scientific) at a 1:20000 dilution. PVDF membranes were visualized by supersignal West Femto chemiluminescence substrate (Thermo Fisher Scientific) and the blot images were acquired using BioSpectrum 810 (UVP).

### Whole cell recordings

Experiments were executed at room temperature (22 °C). The cells plated on poly-D-lysine-coated coverslips (0.1mg/mL) prepared for electrophysiological studies. Whole cell recordings were made with 1-3 MΩ fire-polished recording electrodes. The extracellular solution contained (in mM): 10 HEPES, 140 NaCl, 1 MgCl_2_, and 1 CaCl_2_ (pH=7.4 with NaOH). The intracellular solutions contained (in mM): 10 HEPES, 130 Na gluconate, 10 NaCl, 1 Mg(gluconate)2, and 0.1 EGTA (pH=7.4 with NaOH). Either 300 μM 2-APB or 100 μM capsaicin was dissolved in extracellular solution. Patchmaster was used for data obtainment and analysis. Cells were stimulated every second from −100 to 80 mV in 180 ms.

### Ionomycin calibration

Ionomycin (Thermo Fisher Scientific) was used in calibration for derived calcium concentration from ratios of emitted fluorescence given by 340/380 nm UV excitation. Ten EGTA-buffered standards with different free calcium concentrations were prepared by mixing 0 μM calcium buffer (10mM EGTA in 100mM KCl, 30mM MOPS, pH 7.2) and 39 μM calcium buffer (10mM CaEGTA in 100mM KCl, 30mM MOPS, pH 7.2), the preparation was a serial dilution top-down with Ionomycin (2μM). Cells pretreated with Fura dyes were incubated with each buffer at 30°C and followed by the recording method described under calcium imaging assay section.

## Results

### Capsaicin-elicited influxes of calcium or strontium ions could be monitored in real time

Direct application of 30μM capsaicin evoked saturated cellular ionic fluxes through ligand gated TRPV1. This channel can conduct divalent metal ions from the IIA group (Bouron et al., 2015), among which Ca^2+^ and Sr^2+^. Ion fluxes can be monitored with Fura 2 to emit concentration dependent fluorescence at specific wavelengths. While in physical extracellular calcium concentration (1-2mM Ca^2+^), capsaicin-induced calcium entry has already reached its maximum. By contrast, strontium fluxes were extracellular strontium concentration dependent. 10mM strontium in bath solution caused even higher intracellular strontium concentration than to in 1mM strontium solution (Figure 1). Taken together, fura dyes imaging is a non-invasive technique suitable for monitoring live cells vanilloid responses. Besides, TRPV1 demonstrates higher permeability to Ca^2+^ than to Sr^2+^.

**Figure 1.**
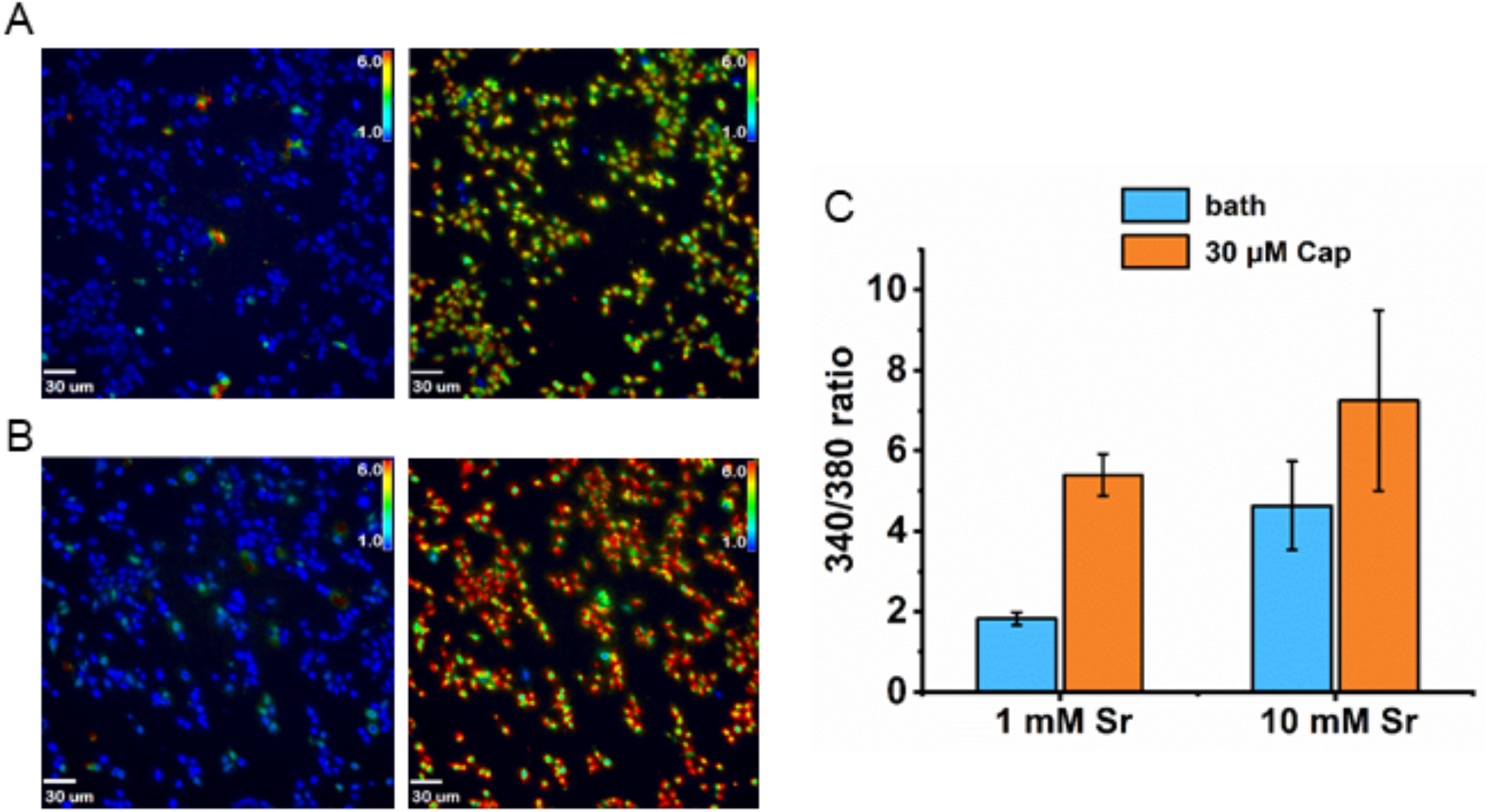
Fura-2 imaging of TRPV1 stably expressing HEK293 cells in strontium solution. (A) extracellular solution contained 1mM Sr^2+^. (B) extracellular solution contained 10mM Sr^2+^. (C) Bar graph showed 30 μM capsaicin induced strontium concentration increase.

Lots of calcium ions flowed into cells due to activation of TRPV1. Non-invasively loading divalent cation sensitive indicators permitted time lapsed Ca^2+^ imaging for continuous activity tracking from a large population of single cells. TRPV1 activity was revealed through a variety of Fura dyes, Fura-2 (Kd = 145nM), Fura-4F (Kd =770 nM) and MagFura-2 (Kd = 25μM) included; they disclosed Ca^2+^ in the entire physiological range for cross-referencing (Table 1). The influx of strontium was significant less than the calcium entry.

**Table 1.**

| **Calcium concentration of capsaicin-induced response detected by different dye (n=3~4 experiments)** |  |
| --- | --- |
| Dye | 30μM Capsaicin application<br>Intracellular [Ca<sup>2+</sup>] (μM) |
| Fura-2 | 6.468±1.888 |
| Fura-4F | 21.004±0.428 |
| Mag Fura-2 | >25 |

The status of Sr^2+^ made dynamics ideal indicator for cells. These data showed that capsaicin induced cation rise were primarily through the TRPV1 channel, not other calcium source. Intracellular calcium concentration can reach at least 25 μM induced by 30 μM capsaicin. Calcium was highly permeable to TRPV1 did serve main divalent ions as the major message carrier in this pathway.

### Mixing in S512F subunit(s) to a receptor specifically reduces capsaicin-evoked channel activity

Each wild-type TRPV1 channel is formed by four identical constituent subunits. It was reported that single Tyr511 residue mutating one subunit of the tandem tetrameric Y511A could maintain full efficacy of the YYYY tandem (Hazan et al., 2015) One capsaicin molecule binding to a tandem tetramer suffices to fully open such TRPV1 channel, suggesting that multiple binding sites could be redundant. We wondered whether a few of those serve as spared subunits just to safeguard the fidelity and robustness of pain transmission.

Y511A must have residual capsaicin sensitivity rather than functioning as capsaicin insensitive (ligand-null) (Figure 2A). We wanted to utilize the most essential residue to do experiments. Therefore, we switched to S512F mutants for further analysis of the minimal number of wild-type subunits required for full agonism of capsaicin receptors. A critical serine residue (Ser512) located at the junction transiting the first intracellular loop into TM3 (3rd trans-membrane segment) was identified a critical residue for capsaicin- or proton-evoked TRPV1 responses (Jordt S.E. and Julius D. 2002).

**Figure 2.**
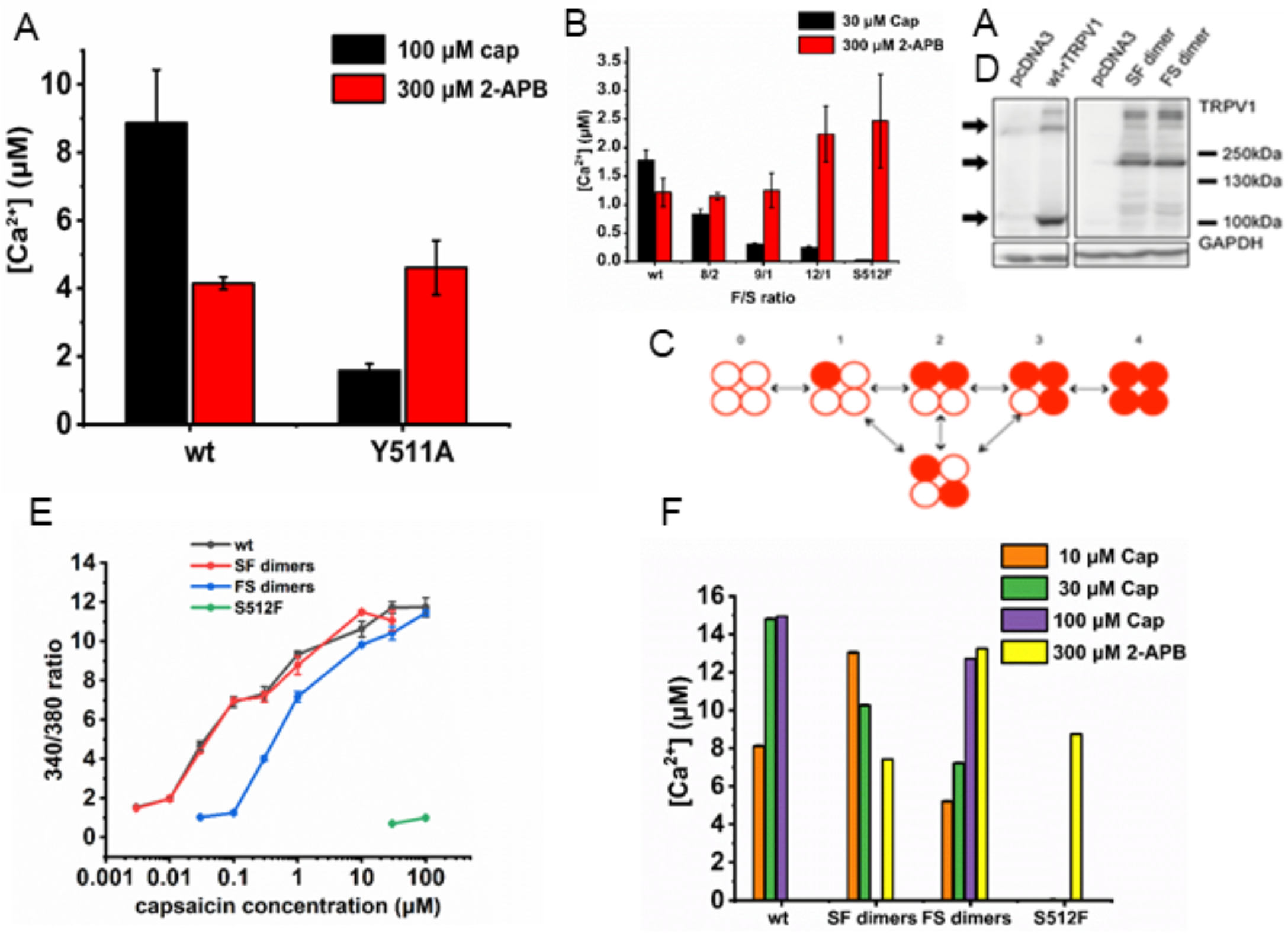
Comparing Ca^2+^ influxes among various receptor constructs (n = 3~4 experiments). (A) Response induced by 2-APB or Cap through activating wild-type trpv1 or Y511A trpv1 (B) Mixing the wild type and the S512F mutant revealed functional dominance of the former. (C) Schematic diagram shows possible constructs are composed of wt-monomer and S512F monomer. (D) Western blotting showed comparable expression of wild-type and mutant channels. (E) Emission ratios of fura-2 loaded cells. (F) The responses were expressed in Ca^2+^ concentrations

We following started from a mixture of wild type and S512F plasmids in a designed ratio expecting to translate and assemble TRPV1s with one wild type plus three mutant subunits to presume full efficacy. Also, we mixed rTRPV1 wild type versus mutant cDNAs in various ratios to create a collection of tetrameric channels to mimic mutants of 0 to 4 capsaicin-bindings stepwise (Figure 2B,C), among which Phe512 was changed back to the wild type Ser512. We found biased mixing in higher ratios of S512F mutants to wild type produced more than simply right shift of capsaicin dose-response curves (potency); rather, capsaicin efficacy was compromised when we only kept one wild-type binding site (Figure 2B).

We initially thought that the repeated use of the same subunits, four non-binding vanilloid insensitive S512F subunits should act as silent receptors. Hence doping a small amount of the wild-type subunits into S512F might markedly rescue non-functional S512F tetramers. Such expression was intended by biased mixing of plasmids. We presumed that the according synthesis could generate functionally conducting tetramers of three S512F plus one wild-type subunits, at the expense of producing plenty of vanilloid non-responding (S512F)4 in the same cell (Figure 2E). We encountered resistance to titration by S512F contrary to this prediction, requiring an extreme ratio bias to titrate away capsaicin evoked Ca^2+^ response according to measured Fura fluorescence. This cast doubts on whether the S512F mutation was truly a general ligand-insensitive receptor or was turned solely capsaicin irresponsive but remained otherwise functional.

To completely probe into the efficacy of the mutant channels, a structurally unrelated non-vanilloid full agonist 2-aminoethoxydiphenyl borate (2-APB) was tested on these constructs (Colton M.X. and Zhu C.K. 2007). The S512F mutant measured no recordable capsaicin induced responses by cytoplasmic Ca^2+^ change. We had similar refractoriness to 2-APB stimulation. To our surprise, 2-APB drove S512F receptors open no less than the capsaicin activated wild-type TRPV1 channels; there is no titrating away the capsaicin unresponsive (S512F)4. Therefore, S512F was functional as long as efficacious ligands are supplied. These findings switched us to pursue covalently linking subunits when creating stoichiometrically fixed tandem receptors comprising of wild type and S512F subunits.

We first made dimeric TRPV1s that might zoom in for comparing the effectiveness of each agonist-bound subunit in contributing to the final response by agonists (Figure 2D). One wild type plus one S512F formed a dimerized receptor cDNA whose expression led to channel opening by capsaicin with full ion conducting efficacy (Figure 2E). A pair of dimeric SF or FS assembled to present a pore lined by two dimerized-dimers, which exhibited the same maximal level of Ca^2+^ entry under 2-APB activation albeit of different subunit permutation orders (Figure 2F). Provided that two of the four of the 6-pass membrane span moieties are wild type, the final receptor complex suffices to encode the full range of pain.

### Tetrameric rTRPV1 requires two or more wild-type subunits for full-range translation of pain

The further determination of the number of subunits required for vanilloid activation of TRPV1 demanded application of 30-100μM capsaicin to fully stimulate individual tetramers and measuring their according maximal activation. The non-S tetramer (FFFF) responded with essentially nothing ([Ca^2+^] = 0.033 μM). The averaged peak response from stimulating the single-S tetramer SFFF was 0.17 μM (Figure 3a, n=3 wells from 162-258 cells). Electrophysiological data also showed that single-S SFFF was not enough to induce full activation when applying 100 μM capsaicin (Figure 3B). Even examined individually, the maximal [Ca^2+^] attainable in any cell expressing mono-S tetramers never had more than 0.5 μM free cytosolic calcium. Contrasting the unrelated pan-TRPV agonist 2-APB, we could confirm comparable receptor expressions; 300 μM 2APB-induced activations were strong and uniform across all tandem tetramers irrespective of the identity of the amino acid residue at the 512 positions. Hence 2-APB does activate the TRPV1 channel by a mechanism quite different from vanilloid induction (Bouron et al., 2015). The binding site in TRPV1 2-APB use is different from capsaicin use. Moreover, all the tetramers were activated by 2-APB to reach comparably high calcium concentrations, supporting comparable protein expression for all single-S tetramers. Double-S or triple-S tetramers both attained full activation at 30 μM capsaicin and their expressions were no less than SSSS. Together, the results suggested that channel activations of TRPV1 were far from the attainable maxima when only one capsaicin-binding site was occupied by vanilloids. A pair of occupied capsaicin bindings sufficed to elevate channel efficacy toward full, bringing the calcium concentrations to match tetramers containing two or more bindings, of which equaled to the maximum given by the wild type.

**Figure 3.**
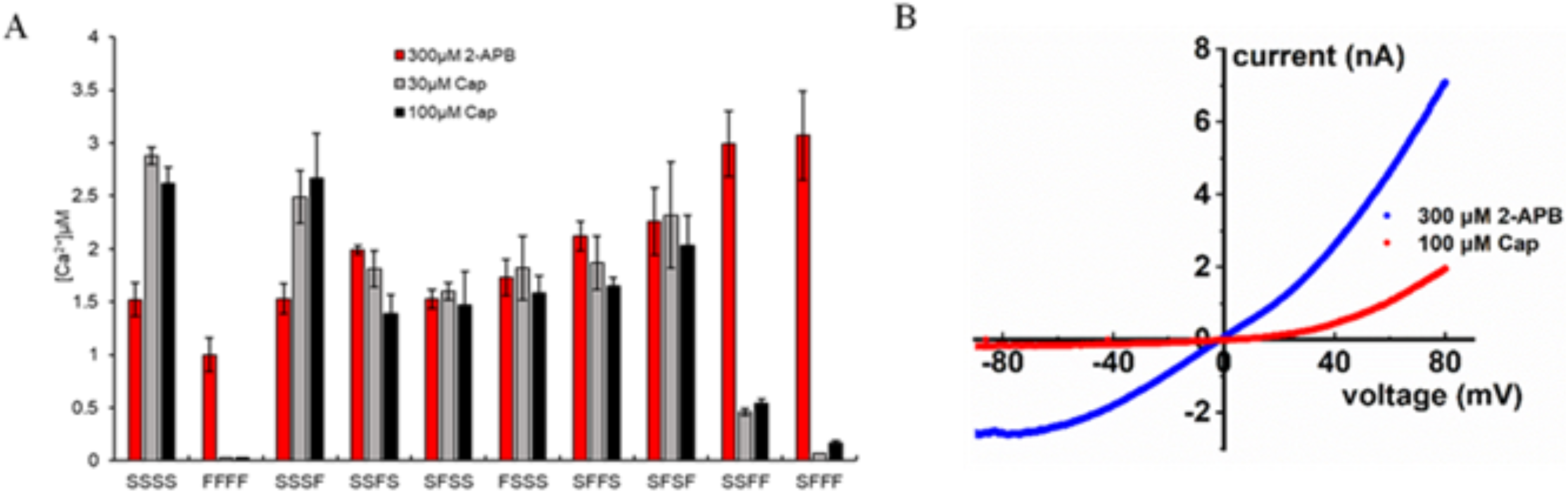
SFFF can’t reach full activation induced by capsaicin. (A) Comparing Ca^2+^ influxes among different tetrameric rTRPV1 constructs (n=3 experiments). (B) voltage-current traces showed apply capsaicin (Cap; 100 μM; red line) or 2-APB (2-APB; 300μM; blue line) to HEK 293t transiently expressing the SFFF construct.

### The partial agonist AEA also requires binding two ligands to exhibit its maximal efficacy

Partial agonist anandamide (AEA) was non-pain-producing. Even administering AEA at its maximal water solubility did not produce a response comparable to capsaicin for the wild-type TRPV1; it did not even reach the calcium level elicited by maximal capsaicin in mutant tandem receptors with just two wild-type binding sites. Agonist effect of AEA was occluded after capsaicin priming (Cao et al., 2013; Ross R.A. 2003). The calcium concentration in SFFF cells ([Ca^2+^] = 0.06 μM) stimulated 30μM AEA by was far less than for quadruple-S ([Ca^2+^] = 0.39 μM) or SFFS ([Ca^2+^] = 0.30 μM) (Figure 4). These data supported that full AEA activation of TRPV1 also required two or more wild-type subunits. Besides, the data showed AEA’s low efficacy wasn’t due to the binding insufficient. The lack of agonist efficacy agreed with the nonpain producing nature of AEA, and vanilloids the pore with the same pharmacological profile.

**Figure 4.**
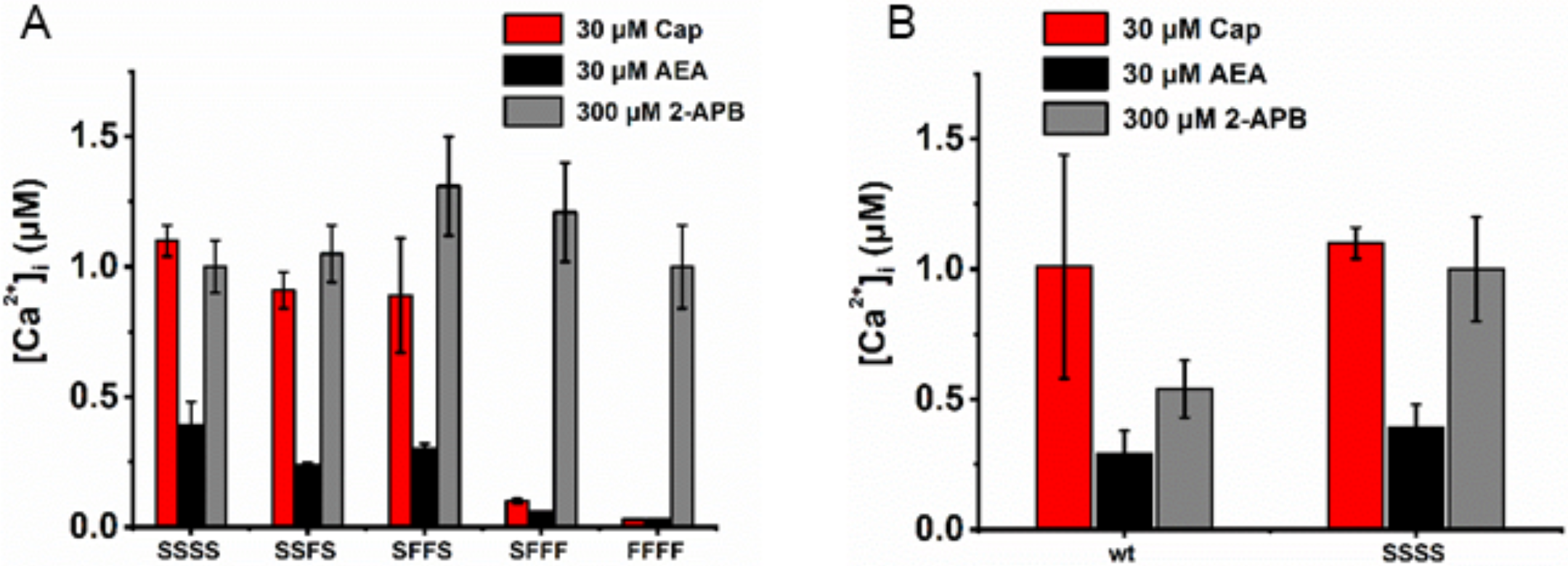
(A) 2-APB is a full agonist compared to capsaicin in the tandem with a 0-4 wild type subunits stepwise, while AEA is a partial agonist throughout. (B) Comparing responses between wild type receptor and tandem-S receptor(SSSS). (n = 3~5 experiments)

## Discussion

Pungent vanilloid compounds isolated from natural source have a wide range of potencies and efficacies, matching well with “spiciness” from a delightful culinary experience to obnoxious chemical deterrence. Capsaicin activation of TRPV1 in expression systems apparently has a threshold far lower than the pungency equivalent indexed by Scoville units. Receptor activation does not necessarily indicate pain; rather, it functions within a wide dynamic range, of which coverage of sensation below conscious awareness. Although pain is indispensable in our daily life, it remains blurred if not impossible to code it to excellent precision. Practically, our analysis projects toward the potential to develop an objective metrics to aid scoring pain modifiers acting through TRPV1, a crucial intermediate step to efficacious and titratable pain medicine.

Signaling pain by an ionotropic receptor can endow a remarkable responding speed and direct additivity. The processing primary input via ion channels follows simple summation, which would be straightforward and effective. It translates inflicts sensibly and speedily enough to alert recipient to awareness of potential or true physical harms in time. A boundary is needed for differentiating the subconscious detection from the sensual awareness of physical pain. TRPV1, the capsaicin receptor, thus serves a critical node dictating bifurcation of nociception. It stands out as a tangible target for modifying pain sensation. TRPV1 activation participates in both the sensory and the motor limbs of organismal responses to pain. TRPV1 functions as a natural device to integrate modes of pro-algesic or painful stimulations. Graded TRPV1 channel related pain reflects resolution of stimulus intensity (Hui et al., 2003). Above all, the understandings about threshold or dynamic range of pain sensation show particular application potentials.

Priel and colleagues had used electrophysiology to show binding of one capsaicin molecule maximally opened a channel, provided that the membrane was held at a sufficiently positive polarization potential. The mutant TRPV1 Y511A had reportedly selectively lost its capsaicin sensitivity in the same electrophysiological studies (Hazan et al., 2015). Due to technical limitation, voltages could only be applied in a range that the membrane could withstand during experiments. To circumvent this barrier, we searched for bath solutions that might augment cation influxes without excessive voltage drive in intact cells. We observed that Y511A remains a vanilloid-sensitive ion channel that responds to capsaicin with reduced affinity. Besides, Y511A activation by heat and/or proton was preserved (Hazan et al., 2015). Other mutations in the neighborhood of Y511 were thus sought for further disruption of vanilloid gating but preservation of activation by non-vanilloid agonists, bringing us to the S512F mutation. Though behaving comparable to the wild-type receptor when challenged with 2-APB, S512F still differs from other species variants of TRPV1 that display drastically lower capsaicin affinity.

At least two of the four subunits need to become ligand bound to evoke full agonist activation in tandem-tetramers, administering capsaicin or AEA on concatenated constructs composed either of capsaicin-insensitive S512F subunit or of the reduced-sensitivity subunit Y511A.

Binding site occupancy commonly drives channel opening to grant influxes of permeant ions in ligand-gated ion channels (Yang et al., 2015; Elokely et al., 2016). A homo-tetrameric TRPV1 with two capsaicin bound subunits has already been bestowed openings enough for raising cytosolic calcium to the maxima not any lower than wild-type receptors that are fully capsaicin-liganded. Pain sensing hence appears built with immense reserves so that it rarely fails. The oxidative sensitization of TRPV1 also demands two cysteine-containing subunits to form the disulfide bond required for the enhancement of channel sensitivity. The “requirement of more than two” underlies a fundamental principle of TRPV1 related pain sensation. We want to point out that the first ligand binding does not bring about enough ion fluxes for reaching pain, presumably securing functional fidelity of chemical nociception to safeguard against an overreaction in case of their accidental encounter. TRPV1 coded nociception covers an ample dynamic range and reserve, demonstrating outstanding economy with its broad conveyance of both painful and innocuous sensations. We measured ligand-induced Ca2+ rise to express stimulus strength dependent activation. Single capsaicin molecule occupancy on a channel yields less than 10% of full efficacy; it would function as a non-painful stimulation unless one can raise activated receptor numbers to unrealistically high overwhelming a cell, a level far higher than our current case. The next level of high response from our single cell studies has already elicited maximal pain, with each dimer-dimer assembled tetrameric channel binds two capsaicin molecules. TRPV1 signaling thus is so robust as to report true pain without fail.

The agonistic activity of capsaicin in TRPV1 gating for both channels display similar normalized dose-response curves for activation. Capsaicin is a lipophilic molecule that binds to TM3 of TRPV1, presumably aided by local lipid interaction provided by the membrane environment (Cao et al., 2013, 2013; Hanson et al., 2015). Bulk capsaicin binding affinities are similar among cells expressing monomeric TRPV1, wild type or mutants, in a bilayer environment. The cation permeation pathway is an ion-mediated structural change in the selectivity filter, and it may generate a higher energy state intermediates with the selectivity filter vacant in transition (Darre et al., 2015). The results from mutant receptors suggested that the principle of mass action might also be exploited by TRPV1 to differentiate pain-producing milieu. Our study could be a lead for guiding finer tuning to find more sensible modifications of molecules or application strategies. Those may be used to counter or even prevent unwanted TRPV1 related

## Author Contributions

TYL, YC, HRM and DSC performed the calcium imaging. TYL performed the western blot. HRM performed electrophysiological experiments. TYL made constructs. TYL, YC, and HRM designed the experiments and wrote and revised the manuscript. HHC designed the experiments, supervised the progression, wrote and revised the manuscript, and communicated with editors

## Acknowledgements

We thank Dr. Hsueh-Chi Yen for stimulating discussion and insightful comments in organization of the finishing draft.

